# Hidden diversity and host specificity of *Cladocopium* symbionts revealed by phylotranscriptomics in giant sea anemones

**DOI:** 10.64898/2026.09.16.749314

**Authors:** Ethan Rickards, Saacnicteh Toledo-Patino, Malgorzata Hall, Saori Miura, Matthias Wolf, Bruno Frédérich, Konstantin Khalturin, Vincent Laudet

## Abstract

Giant sea anemones form an intricate, metabolically linked symbiosis with anemonefish (genus *Amphiprion*) and dinoflagellates (family Symbiodiniaceae). While the association between fish and anemone is well documented, the identity and distribution of resident Symbiodiniaceae remains poorly characterized. Here, we used RNA-sequencing of anemone tentacles and post-assembly filtering to recover Symbiodiniaceae sequences from Japanese anemones. Using transcriptome-wide phylogenetic approaches, we identified distinct Symbiodiniaceae strains in seven sea anemone species and four cryptic lineages of *Entacmaea quadricolor*. Phylogenies of 1300 and 214 nuclear genes resolved relationships among these symbionts with greater sensitivity than traditional molecular markers such as the Internal Transcribed Spacer 2 (ITS2) or partial 28S ribosomal DNA. Marker-based assessments indicate these Symbiodiniaceae as novel and specific to sea anemone hosts. Our transcriptome-wide phylogenies reveal previously undescribed *Cladocopium* (Clade C) strains associated with giant sea anemones, with symbionts clustering according to host species and lineage. Cophylogenetic analysis revealed significant phylogenetic congruence between anemone and Symbiodiniaceae, indicating a nonrandom association between host and symbiont evolutionary histories. Two sympatric cryptic lineages of *E. quadricolor* previously identified through host transcriptomics harbor distinct, specific Symbiodiniaceae assemblages. Analysis of genes under positive selection indicates functional divergence between these Symbiodiniaceae; three-dimensional reconstructions of symbiont cells using serial block-face scanning electron microscopy show consistent differences in cellular organization between two strains, further supporting their distinct functional identity. Together, our results uncover a hidden diversity of host-specific Symbiodiniaceae and suggest that cryptic diversification in giant sea anemones is mirrored by diversification of their algal symbionts within this tripartite symbiosis.

## Introduction

The relationship between anemonefish (genus *Amphiprion*) and giant sea anemones is a classic example of mutualistic symbiosis. Similar to Scleractinia (reef-building corals), giant sea anemones host photosynthetic dinoflagellates belonging to family Symbiodiniaceae (zooxanthellae) within their tentacles. Symbiodiniaceae are involved in numerous marine symbioses, and the disruption of this association represents a critical area of research [1,2]. In this relationship, Symbiodiniaceae impart photosynthetic byproducts to their host in exchange for a protected, light-rich habitat abundant in nutrients such as nitrogen [3]. These relationships are essential, as photosynthetically derived carbon supports host growth, maintenance, or daily function [3,4]. In context of the fish-anemone-dinoflagellate symbiosis, anemonefish strengthen this exchange by excreting a copious amount of ammonium that is assimilated by the giant sea anemone [5]. Studies on non-fish-hosting sea anemones show a host-driven limitation of inorganic nitrogen to directly regulate Symbiodiniaceae densities [6]. As a consequence, anemones support a greater density of Symbiodiniaceae when associated with anemonefish [5,7]. These demonstrate the ongoing reciprocal nutrient exchange between all three partners and underscore the physiological integration of this tripartite symbiosis [8,9]. Given its complexity, understanding this intricate relationship between anemonefish, giant sea anemones, and Symbiodiniaceae requires characterization of each partner.

This symbiosis involves approximately 29 anemonefish and 10 giant sea anemone species, the latter of which encompasses three distinct morphotypes and genera: carpeting (*Stichodactyla*), long-tentacle (*Heteractis* = *Radianthus*), and bubble-tip (*Entacmaea*) sea anemones [10–14]. The associations between fish and anemone are non-random and display varying degrees of specificity, with some anemonefish species acting as strict specialists restricted to a single host (e.g. *A. frenatus* with only *E. quadricolor*), whereas generalists are capable of associating with multiple anemone species (e.g. *A. clarkii* with 5 natural hosts in Okinawa) [15–18]. This variability plays an important role in ecological differentiation and species coexistence within reef communities [18–20].

However, this general framework underestimates the true complexity of these associations, as recent studies suggest previously unrecognized cryptic diversity within a single anemone species, *Entacmaea quadricolor*. In Japan, *E. quadricolor* comprises four cryptic lineages (A–D), two of which coexist in sympatry in Okinawa (A and D) [21,22]. Although morphologically similar, these lineages are genetically distinct and associate with different anemonefish species: *A. clarkii* preferentially associates with lineage A, whereas *A. frenatus* inhabits lineage D [21,22]. This indicates a cryptic diversity of sea anemones, each with potentially distinct symbiotic relationships. Though, whether such diversity is mirrored in their Symbiodiniaceae endosymbionts is unknown.

Despite advances in our understanding of the anemone-anemonefish symbioses, the distribution of their resident Symbiodiniaceae remains unclear. Symbiodiniaceae are symbiotic with a wide range of marine hosts, and these associations also display variable degrees of specificity [23]. Some are restricted to a narrow range of hosts, whereas others are generalists capable of associating with multiple species [24–27]. Further, some hosts may also harbor multiple species of Symbiodiniaceae at a time [24,28]. As zooxanthellae species may vary in thermal, irradiance, or nutritional preferences, their identity is important to assess the functionality and stability of the symbiotic relationship [29–32].

Despite their importance, the Symbiodiniaceae partner within this tripartite symbiosis has not been well-characterized. The few studies that do assess this symbiosis examine only one host species in a specific area and largely indicate them as hosting *Cladocopium* (formerly *Symbiodinium* Clade C), a genus widespread in Indo-Pacific hosts [31,33–36], and rarely multihosting with *Symbiodinium* (Clade A) [37]. In Japan, it is known that giant sea anemones host *Cladocopium*, though little else about this symbiotic relationship is known [13].

These studies implement molecular markers, namely the multi-copy internal transcribed spacer 2 (ITS2) and 28S Large Ribosomal Subunit (LSU) rDNA. Due to ITS2’s ease of amplification and historical precedent, this 200-bp marker is the most used method to identify Symbiodiniaceae [23,38,39]. However, distinct species are known to share identical ITS2 sequences, obfuscating algal identity when relying solely on this marker [23,27,40]. ITS2 is also prone to intragenomic variation, with a single species possessing multiple variations of ITS2, making it difficult to discern species-specific differences [38]. In contrast, LSU is a 620-bp sequence that has been used in conjunction with ITS2 and other marker sequences to redefine the nine current subclades of Symbiodiniaceae [40]. While also multi-copy, LSU sequences are more conserved, but this lends itself to LSU failing to resolve between species within a clade [23,40]. These issues amongst molecular markers make them unreliable benchmarks to accurately differentiate Symbiodiniaceae further than the genus level.

Whole genome sequencing has been used to distinguish Symbiodiniaceae with a higher sensitivity than marker-based approaches while identifying key genetic differences on a population level [41]. These Symbiodiniaceae genomes can be expansive and complex, with some reaching up to 5 Gbp [42]. As such, transcriptome-wide phylogenies may detect Symbiodiniaceae at a high sensitivity while avoiding the limitations of marker or full genome sequencing. Phylotranscriptomics targets a subset of the genome, while being sufficient to accurately describe and differentiate between samples [43]. As these genes are all expressed, further analysis into the transcriptome can also provide insight into functional differences and biological processes between Symbiodiniaceae [44]. This approach has been successfully conducted for sea anemone phylogeny, but not yet for their hosted Symbiodiniaceae [13,21].

Here, we aim to characterize the missing diversity and distribution of Symbiodiniaceae associated with Japanese giant sea anemones. Using transcriptome-wide phylogenetic analysis across seven of the ten anemone species (*E. quadricolor*, *Heteractis aurora*, *R. crispa*, *R. magnifica*, *Stichodactyla gigantea*, and *S. mertensii*) and all four *E. quadricolor* cryptic lineages, we test whether Symbiodiniaceae exhibit specialization and patterns of codiversification with host anemones. We then examine whether phylogenetic divergence is associated with functional differences through further analysis of positively selected genes and three-dimensional ultrastructural comparisons of those belonging to sympatric *E. quadricolor* lineages. These findings provide a foundation for understanding the diversity, distribution, and evolutionary history of Symbiodiniaceae within this iconic tripartite symbiosis.

## Results

### LSU and ITS2 markers identify all symbionts as *Cladocopium* but provide limited phylogenetic resolution

Fifty-one giant sea anemones representing seven species (*Entacmaea quadricolor, Heteractis aurora*, *Radianthus crispa*, *R. magnifica*, *Stichodactyla gigantea*, *S. hadonni* and *S. mertensii*) and four cryptic lineages of *Entacmaea quadricolor* were analyzed for their associated Symbiodiniaceae communities.

We identified Symbiodiniaceae according to previous studies relying on the molecular markers LSU or ITS2. Marker sequences for both LSU and ITS2 were extracted from assembled transcriptomes via blastn (1e-20) (45) and compared to curated marker databases. LSU sequences were found in 49/51 transcriptomes, yielding 35 unique LSU sequences (Supp. Materials 1). These sequences had 0-19 base pair differences from any of the references. Complete ITS2 sequences were found in 39/51 of the transcriptomes. 24 novel ITS2 sequences (Supp. Materials 2) were isolated, with 1-53 base pair differences from the references. Both LSU and ITS2 sequences consistently assigned sequences within genus Cladocopium. As these are largely novel sequences, samples are named according to their genus and host anemone species.

Extracted Symbiodiniaceae LSU and ITS2 sequences were concatenated to construct a maximum-likelihood phylogenetic tree and assess relationships among samples (Supp. Figure 1). The resulting tree showed limited resolution between Symbiodiniaceae, and the relationships between distinct taxa remain poorly resolved. Many endosymbionts appeared non-monophyletic and distributed throughout the phylogeny. Although this pattern could suggest high within-host symbiont diversity, it is likely an artifact of the multi-copy nature of these markers, which can overestimate Symbiodiniaceae diversity (23, 40). These findings support previous claims that LSU and ITS2 markers provide reliable genus-level identification but are insufficient to resolve fine-scale symbiont diversity.

**Figure 1.**
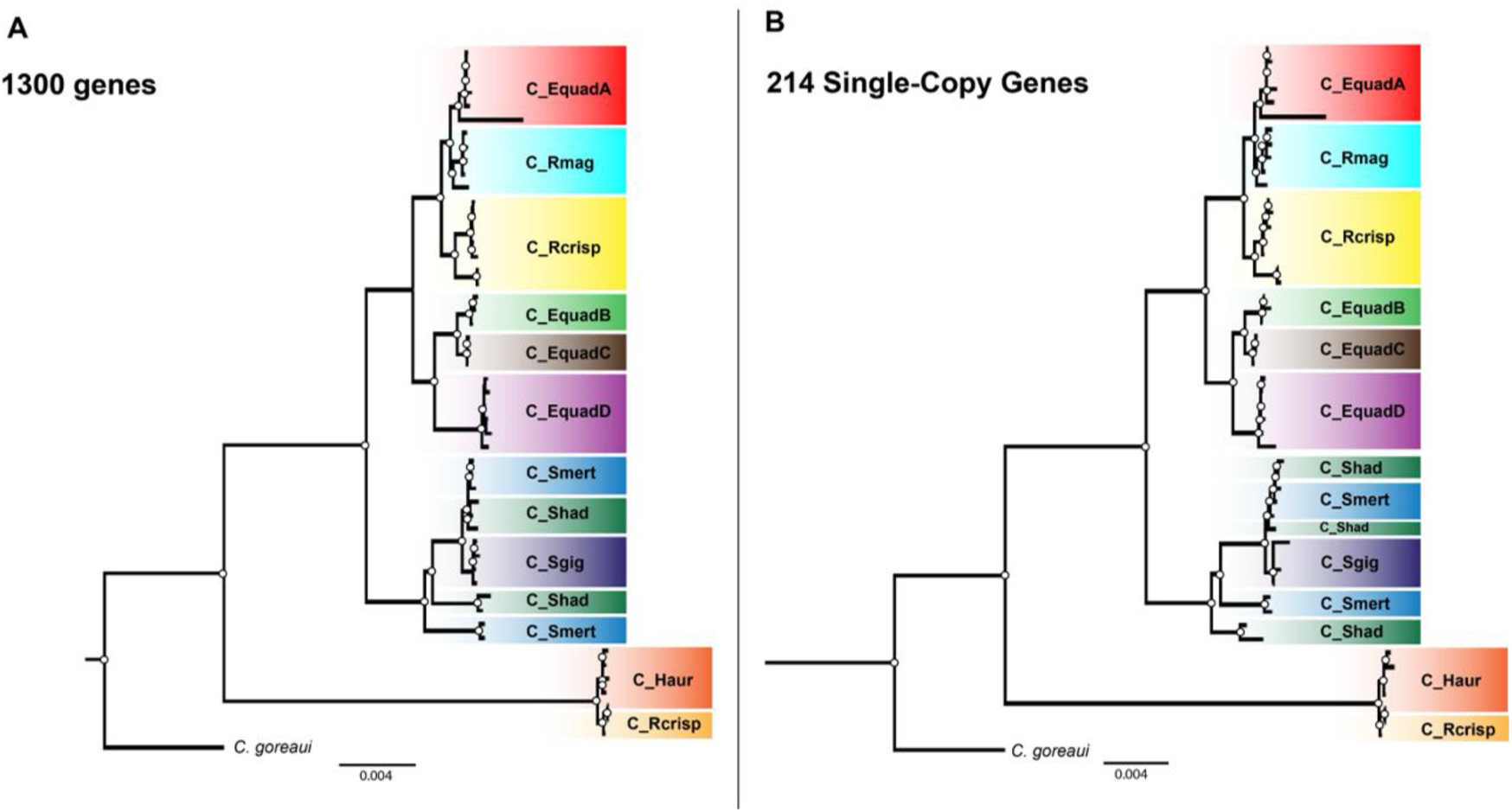
Maximum-Likelihood (ML) phylogenies of Symbiodiniaceae associated with Japanese giant sea anemones. Each sample is color-coded and labeled according to its host anemone. Nodes with bootstrap support >80% (n = 10,000 replicates) are denoted by dots. **A.** ML phylogeny constructed from 1,300 orthologous genes selected using OrthoFinder. **B.** ML phylogeny constructed from 214 single-copy genes.

### Phylogenomics uncovers strong host-associated structuring of Symbiodiniaceae communities

Given the weak resolution of marker-based methods, we implemented multilocus phylogenies based on transcriptomic data to generate more resolutive and robust trees.

A phylogenomic tree constructed from 1300 orthologous genes recovered well-supported relationships among Symbiodiniaceae associated with giant sea anemones (**Figure 1A**). In contrast to the poor resolution obtained with LSU and ITS2 markers, samples are easily distinguishable and the relationships between them are clear. To assess the robustness of these relationships, we generated a second phylogeny using a more stringent choice of 214 strictly single copy orthologs present in all samples (**Figure 1B**). Despite the substantial reduction in gene number, the resulting topology was highly similar to that recovered from the 1300-gene dataset, with comparable clustering patterns. Together, these results demonstrate that phylotranscriptomic analyses provide a robust and highly resolved framework for investigating Symbiodiniaceae relationships.

Visual comparison between the host and symbiont phylogenies show parallel structure between host and symbiont trees, with Symbiodiniaceae clustering according to host anemone species or lineages (**Supp. Figure 2**). Symbionts from the same anemone species or *E. quadricolor* lineage consistently cluster according to host, suggesting a high degree of host specificity. However, the two trees are not fully congruent, as symbionts associated with *E. quadricolor* lineage A formed a distinct clade sister to those hosted by *R. magnifica*, whereas symbionts from lineages B–D clustered separately. Likewise, symbionts associated with *H. aurora* and two *R. crispa* formed a well-supported clade distinct from all other host-associated lineages

**Figure 2.**
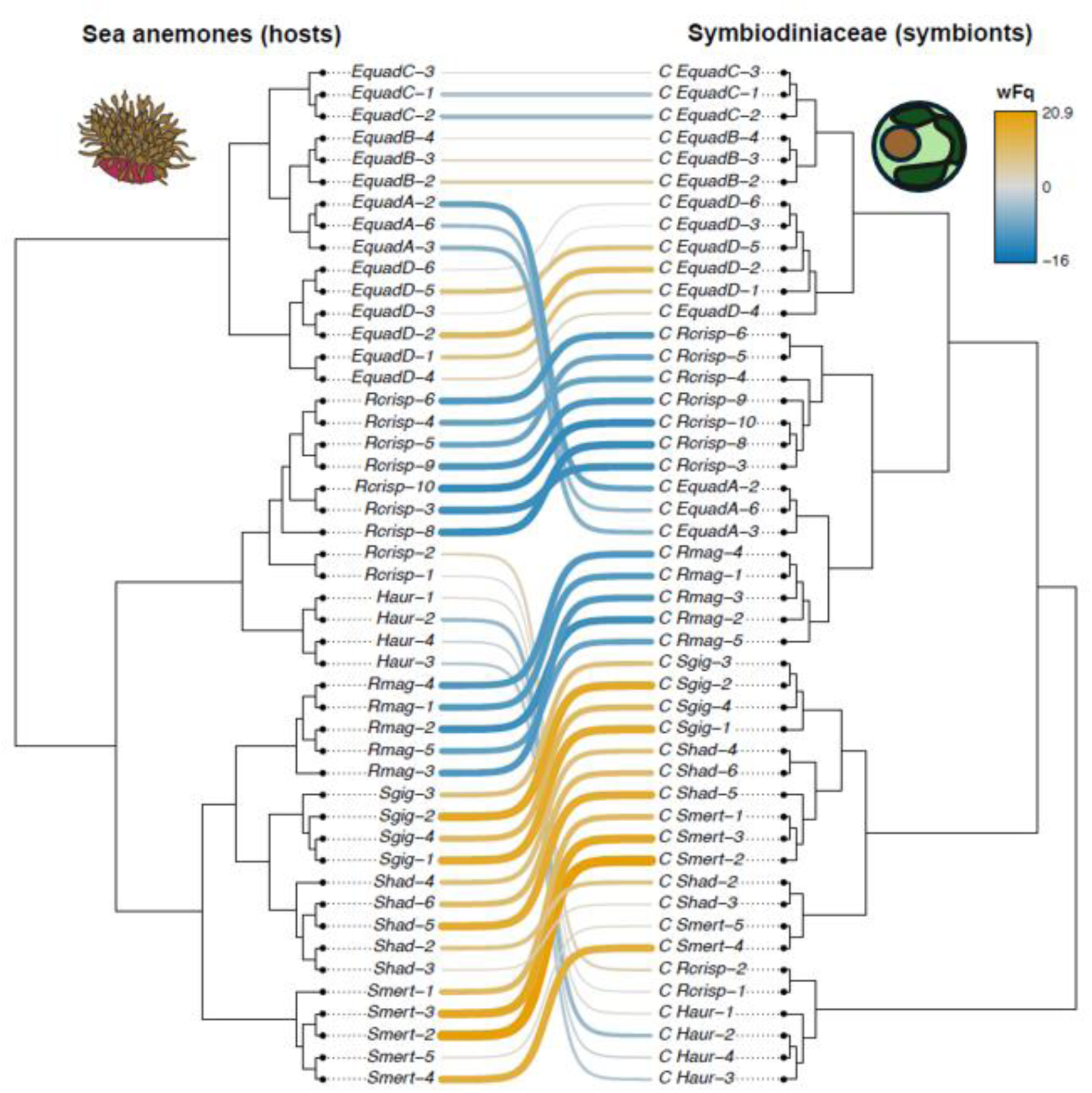
Cophylogram of sea anemone hosts (left) and Symbiodiniaceae (right). Weighted frequencies of host-symbiont associations are determined through Random TaPas are denoted using a color scale from lowest (blue) to highest (orange).

This visual assessment of Symbiodiniaceae and host anemone cophylogeny is supported statistically. We estimated cophylogenetic signal using PACo (Procrustean Approach to Cophylogeny), which evaluates the concordance between host and symbiont phylogenies through Procrustes superimposition of principle coordinate representations of phylogenetic distance matrices [46]. PACo analysis revealed a significant global cophylogenetic signal between hosts and symbionts (SS = 0.63, p < 0.001), indicating that the observed associations were more phylogenetically congruent than expected under random associations. These patterns and cophylogenetic signal are evidence of codiversification between host anemone and Symbiodiniaceae.

We then used Random Tanglegram Partitions (Random TaPas) [47] to decompose the global cophylogenetic signal and identify the strongest Symbiodiniaceae-anemone associations most responsible for the congruence. Random TaPas partitioned global congruence by evaluating the frequency with which individual associations occurred among the most congruent random partial tanglegrams. Here, weighted frequences (wFq) varied widely among interactions (range:-15.99 to 20.92), indicating heterogeneity in the contribution of individual associations to the observed cophylogenetic pattern. **Figure 2** summarizes the relative contribution of each host-symbiont association to the cophylogenetic signal inferred across all randomizations. The associations within the *Stichodactyla* genus showed strong positive wFq values and therefore contributed substantially to the global cophylogenetic signal, whereas the associations within the *Radianthus* genus with negative wFq values contributed comparatively little. This disparity is reflected through phylogenetic positioning; *Stichodactyla* is monophyletic in both host and symbiont, *Radianthus* is monophyletic only in the host. Likewise, *E. quadricolor* lineage A symbionts have a lower wFq compared to the other three lineages.

### Sympatric *E. quadricolor* lineages harbor distinct and evolutionarily divergent Symbiodiniaceae

The distribution of Symbiodiniaceae adds to the ecologically fascinating system of Okinawan *E. quadricolor*. Symbiodiniaceae belonging to *E. quadricolor* lineage A form a sister clade to those hosted in *R. magnifica,* and paraphyletic to those in lineages B-D. Host *E. quadricolor* lineages A and D are sympatric in Okinawa and associated with different species of anemonefish, respectively *A. clarkii* and *A. frenatus*. Their resident Symbiodiniaceae are likewise genetically distinct, which reinforces the notion of different cryptic species of *E. quadricolor*.

The availability of transcriptome-scale data allowed us to move beyond phylogenetic reconstruction and directly investigate functional divergence between Symbiodiniaceae lineages. Given the phylogenetic distance between the symbionts associated with *E. quadricolor* lineages A and D, we tested whether these symbionts show evidence of lineage-specific selection. Therefore, PosiGene [48] was used to identify positively selected genes in C_EquadA *versus* C_EquadD. Analyses of these two groups identified 54 genes under positive selection in C_EquadA and 39 in C_EquadD. Functional annotation was inferred from sequence similarity searches using BLASTP against multiple reference databases, including NCBI nr, *Arabidopsis*, *Homo sapiens*, and Cnidarian protein datasets [49]. Functional domains containing lineage-specific substitutions were identified to highlight candidate proteins that may have diverged between C_EquadA and C_EquadD. Despite the limited annotation of Symbiodiniaceae, several functional categories were identified.

Both C_EquadA and C_EquadD contained positively selected genes associated with cell signaling, transport and trafficking, protein homeostasis, and gene expression (**Supp. Table 1**). Genes related to cell structure and motility were identified only in C_EquadA. Interestingly, genes associated with membrane transport and carbon metabolism were more prevalent in C_EquadD. Conversely, genes involved in amino acid metabolism were greater in C_EquadA.

The 93 identified genes contained amino acid substitutions and/or insertions and deletions that were consistent within each lineage. Several of these changes were in conserved protein domains, suggesting a potential impact on the function of the proteins involved. **Table 1** shows a representative subsection of these genes, their length, predicted function, functional category, domain architecture, and average percent identity in *C. goreaui, D. trenchii*, C_EquadA, and C_EquadD (See **Supp. Table 1** for more). **Figure 3** highlights representative alignments of two positively selected genes per lineage, focusing on annotated functional domains. Alignment for all genes can be accessed in **Supp. Table 1**.

**Figure 3.**
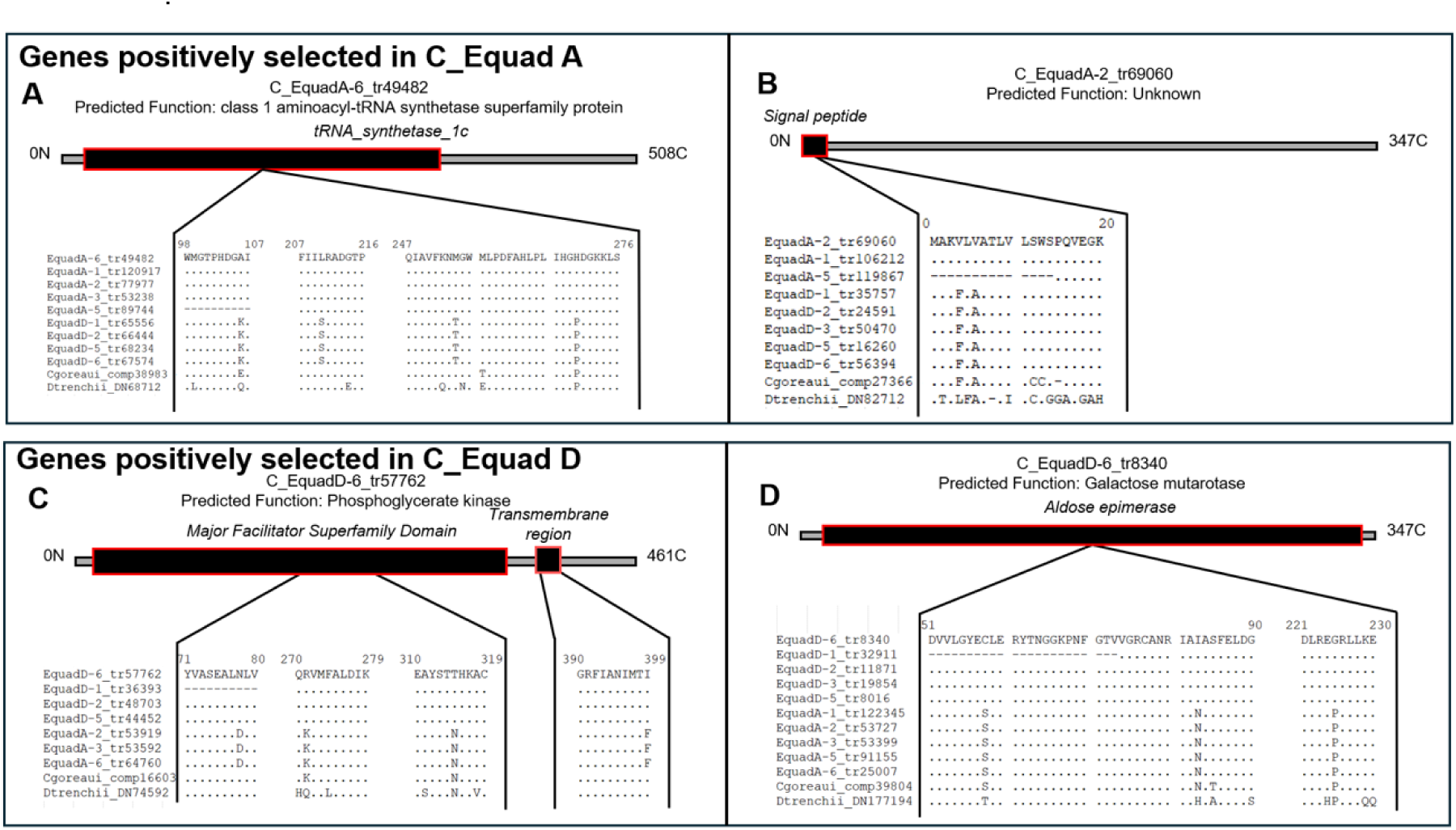
Alignments of functionally important protein domains of Symbiodiniaceae belonging to E. quadricolor A (A, B) and D (C,D) to C. goreaui and D. trenchii. Protein Domains were identified using SMART, and alignments visualized via SeaView. Consistent mutations are observed between Symbiodiniaceae samples, following assemblage lines.

**Table 1.** Positively Selected Genes of *Cladocopium* within *E. quadricolor* lineages.

Four positively selected genes per host lineage were chosen at random.
| Sequence | Sequence Length | Predicted Gene (BLASTp and InterProScan) | Functional Category | Domains with a substitution | *C. goreau* %ID | *D. trenchii* %ID | Average C_Equad-A %ID (Number of Sequences) | Average C_Equad-D %ID (Number of Sequences) |
| --- | --- | --- | --- | --- | --- | --- | --- | --- |
| C_EquadA-2_tr69060 | 799 | Unknown |  | signal peptide | 96.12 | 77.736 | 100 (5) | 98.87 (5) |
| C_EquadA-3_tr64926 | 564 | Threonine dehydratase | Amino acid and cofactor Metabolism | PALP (pyridoxal phosphate dependent enzyme) domain \| Thr dehydratase | 95.567 | 84.041 | 99.87 (4) | 98.40 (3) |
| C_EquadA-3_tr7189 | 525 | Voltage-gated sodium channel | Cell signaling | Ion transporter | 97.333 | 82.634 | 100 (5) | 99.24 (5) |
| C_EquadA-6_tr49482 | 508 | class 1 aminoacyl-tRNA synthetase superfamily protein | Gene expression | tRNA_synt_1c | 96.844 | 82.353 | 100 (5) | 99.01 (4) |
| C_EquadD-1_tr10104 | 548 | Unknown |  | signal peptide | 92.674 | 83.029 | 96.90 (4) | 100 (5) |
| C_EquadD-5_tr46002 | 628 | ABC transporter | Transport and trafficking | ABC membrane transport \| AAA ATPase | 96.513 | 84.227 | 99.04 (3) | 100 (2) |
| C_EquadD-6_tr57762 | 461 | Phosphoglycerate kinase | Carbon and energy metabolism | MFS \| transmembrane region | 98.478 | 84.165 | 98.91 (3) | 100 (4) |
| C_EquadD-6_tr8340 | 347 | Galactose mutarotase | Carbon and energy metabolism | Aldose_epim | 94.767 | 82.647 | 98.17 (5) | 99.11 (5) |

Together, these results indicate that the divergence between C_EquadA and C_EquadD is not limited to phylogenetic differentiation but is also accompanied by functional shifts detectable at the transcriptome level.

### Distinct cellular organization distinguishes *Cladocopium* associated with sympatric *E. quadricolor* lineages

To determine whether the strong phylogenetic divergence observed between the sympatric *Cladocopium* lineages associated with *E. quadricolor* is accompanied by structural differentiation, we reconstructed individual symbiont cells in three dimensions using serial block-face scanning electron microscopy (SBF-SEM). This approach allowed quantitative comparisons of organelle organization and cellular architecture between C_EquadA and C_EquadD. Accumulation bodies, chloroplasts, chromatin, lipid droplets, mitochondria, nuclei, pyrenoids, and starch bodies were each labeled and assessed. Chromatin bodies were labeled when distinguishable, though could not be reliably identified in all cells and were therefore excluded from comparative analyses.

Three-dimensional models of C_EquadA and C_EquadD Symbiodiniaceae cells were constructed to assess morphology (**Figure 4**). Reconstructions revealed that both *Cladocopium* lineages share the archetypal Symbiodiniaceae cellular architecture, with peripheral chloroplast lobes, centrally positioned pyrenoids connected to surrounding chloroplasts by 2–3 stalks, and abundant starch bodies and lipid droplets distributed throughout the cytoplasm. Despite this overall similarity, the two lineages differed in cell size, with C_EquadD cells being consistently larger than C_EquadA cells (448.9 ± 80.2 µm³ versus 341.4 ± 41.2 µm³).

**Figure 4.**
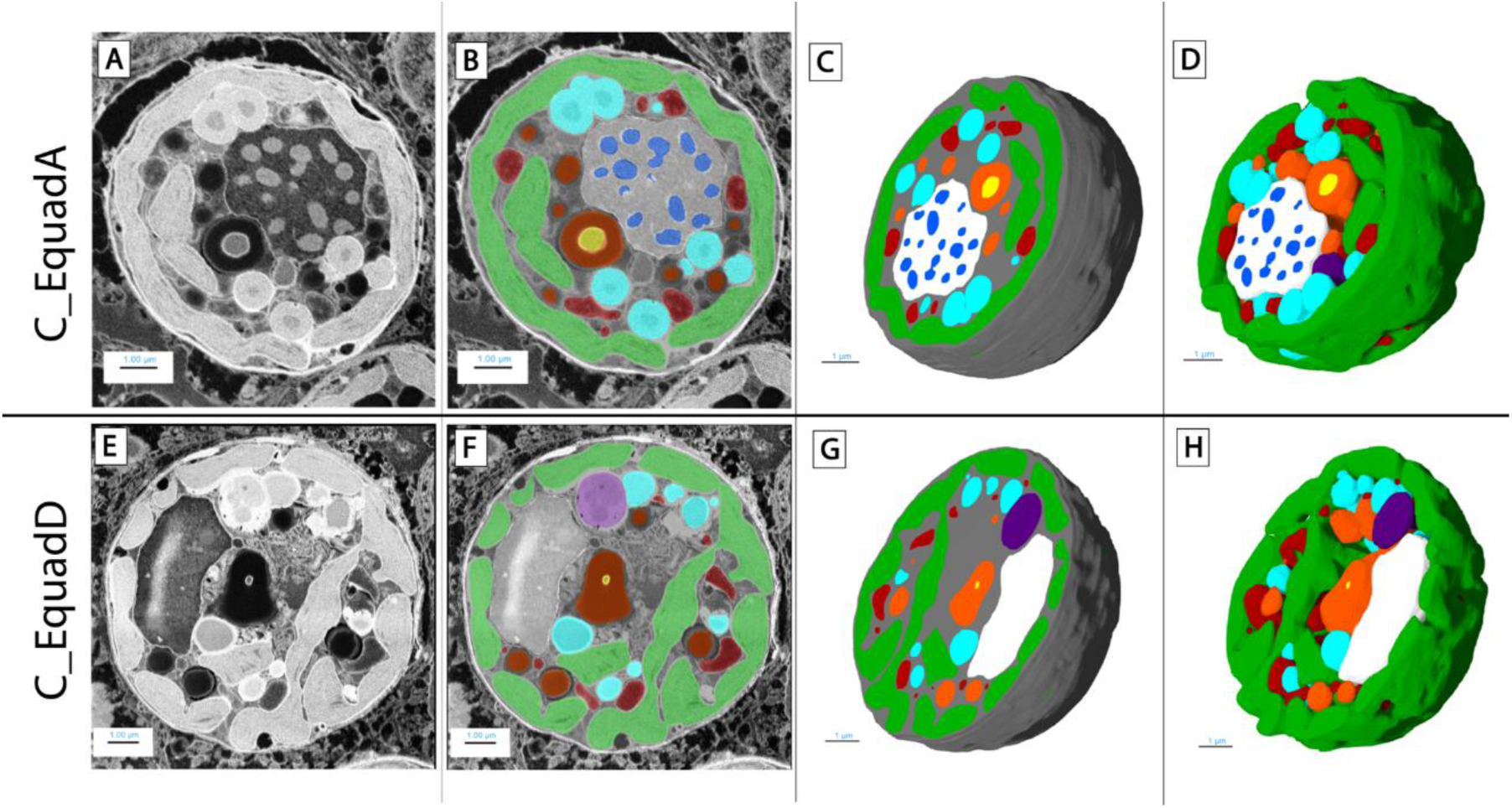
3D models and reconstructions of Symbiodiniaceae cells belonging to E. quadricolor lineage A **(A-D)** and lineage D **(E-H).** The following structures were labeled: chloroplast (green), nucleus (white), chromatin (blue), mitochondria (red), starch bodies (orange), lipid droplet (light blue), pyrenoid interior (yellow), and accumulation body (purple). Chromatin was only labeled in samples where they were discernable in the FIB-SEM data. Unlabeled sections of the cell are gray. **A** and **E** show completed volume reconstruction; **B** and **F** are unlabeled 3D slices of Symbiodiniaceae cells; **C** and **G** are labeled cells. **D** and **H** are 3D reconstructions depicting only the labeled portions of the cell.

To move beyond qualitative observations and obtain quantitative measures of cellular organization, we calculated the volume occupied by each major organelle and expressed it as a proportion of total cell volume (**Figure 5**). This analysis revealed consistent differences between the two *Cladocopium* lineages. Most notably, C_EquadD cells allocated a substantially greater fraction of their volume to chloroplasts (45.37 ± 0.47%) than C_EquadA cells (38.43 ± 1.85%), whereas C_EquadA cells contained approximately twice the proportion of lipid droplets (5.37 ± 1.49% versus 2.60 ± 0.75%). Additional differences were observed in the fraction of unlabeled cellular material. These quantitative observations complement the phylogenomic and positive-selection analyses and indicate that the divergence between C_EquadA and C_EquadD extends beyond sequence variation to encompass differences in cellular organization and physiology. Together, these independent lines of evidence are consistent with C_EquadA and C_EquadD representing distinct *Cladocopium* species.

**Figure 5.**
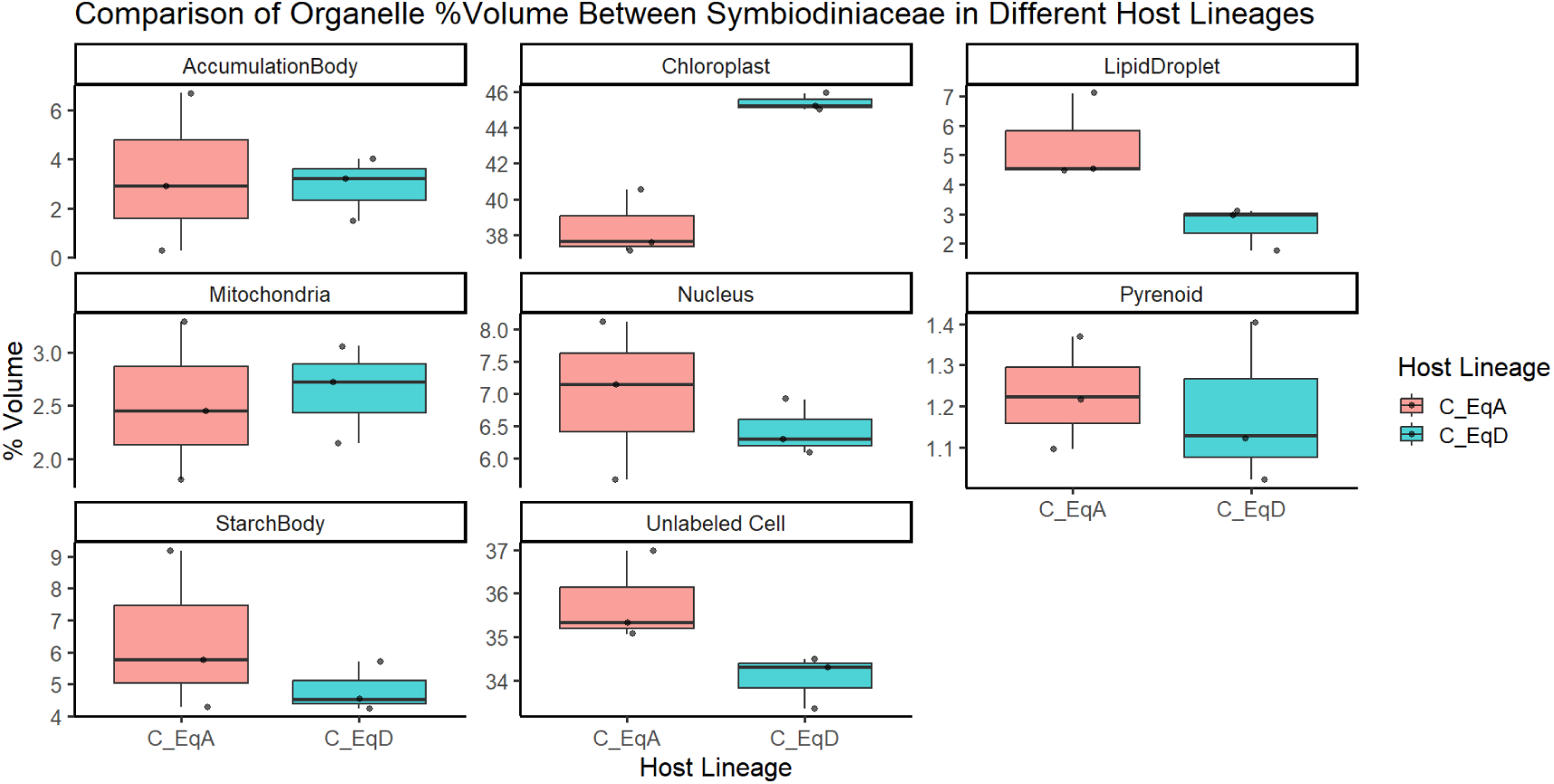
Boxplots of organelle percentage by volume between Symbiodiniaceae hosted by E. quadricolor lineages A (red) and D (blue). Volumes of each label were collected then divided by total cell volume before plotting.

## Discussion

In this study, we combined transcriptome-wide phylogeny, cophylogenetic assessment, analyses of positive selection, and three-dimensional ultrastructural reconstructions to investigate the diversity and evolution of Symbiodiniaceae associated with giant sea anemones. Our results reveal previously undescribed host-specific *Cladocopium* lineages and show a strong congruence of host and symbiont phylogenies. We also found evidence for functional divergence among symbionts using phylogenetic and cellular datasets. Together, these results uncover a hidden level of diversity within the iconic sea anemone–anemonefish–Symbiodiniaceae symbiosis and that diversification of giant sea anemones was mirrored by their algal symbionts. More generally, our study highlights that symbiont diversity in these systems has been underestimated and that transcriptome-wide approaches can reveal patterns of host specificity and divergence that are not detectable using traditional molecular markers alone.

### Phylotranscriptomics provides a robust framework to characterize Symbiodiniaceae diversity

Our results illustrate the limitations of traditional approaches based on ITS2 and LSU for characterizing Symbiodiniaceae diversity. Although these markers allow for reliable assignment of symbionts to the genus *Cladocopium*, they provide little phylogenetic resolution and do not permit the distinction of host-specific lineages revealed by our transcriptomic analyses. This limitation is not surprising, as ITS2 is a multicopy marker subject to substantial intragenomic variation, whereas LSU is generally too conserved to resolve closely related lineages [23,38,40]. In addition, none of the ITS2 sequences recovered in this study were represented in the comprehensive SymPortal database [38] or in the dataset of Shi et al. 2021 [39], further suggesting that these symbionts represent previously uncharacterized diversity.

In contrast, phylogenies reconstructed from hundreds of orthologous genes produced highly resolved and robust trees [13,43], revealing Symbiodiniaceae lineages associated with specific host species and cryptic lineages. These highlight a previously unrecognized cryptic diversity of *Cladocopium* associated with Japanese giant sea anemones. Previous studies based on ITS2 markers identified giant sea anemones as hosting mainly *Cladocopium*, and in one case *Symbiodinium* [34–37]. Likewise, the studied Japanese samples consistently harbored a single host-associated *Cladocopium* lineage per sample, though demonstrate a clear pattern of host-specific association not represented in the above studies. Because geography and latitude are known to influence Symbiodiniaceae assemblages [50,51], some of the differences between this and other studies may also reflect the biogeographic structuring of symbiont diversity.

This approach, however, relies on an important assumption: each anemone contains a single Symbiodiniaceae lineage. If multiple closely related species or lineages were present within the same individual, transcriptome assembly would generate a composite dataset containing sequences from different biological entities, making phylogenetic inference difficult to interpret. Our data suggests otherwise, with the anemones hosting one strain of *Cladocopium*. First, symbionts consistently cluster according to host species and cryptic host lineages. If each sample contained varying proportions of multiple Symbiodiniaceae strains, the resulting phylogenies would be expected to show unstable and incoherent topologies. Second, phylogenies reconstructed from 1300 orthologous genes and 214 strictly single-copy orthologs selected using highly conservative criteria, produced nearly identical topologies. This strong congruence provides additional support for the robustness of our approach and suggests that each sample is dominated by a single biological entity.

More generally, our results suggest that transcriptome-based phylogenomic approaches represent a particularly powerful tool for studying Symbiodiniaceae diversity, host specificity and evolution. They offer substantially higher resolution than traditional molecular markers while remaining considerably more accessible than whole-genome sequencing approaches. We therefore expect phylotranscriptomics to become an increasingly useful framework for resolving cryptic Symbiodiniaceae diversity and investigating patterns of host–symbiont evolution.

### A hidden diversity of host-specific *Cladocopium*

The distribution of *Cladocopium* in Japanese giant sea anemones is not random, but instead strongly structured according to host identity. This study and Kashimoto et al. 2024 [21] utilize different subsets of the same assembled transcriptome, allowing for tandem examination of Symbiodiniaceae and host anemone. Across constructed multi-locus phylogenies, *Cladocopium* clustered according to host species, including the cryptic lineages of *E. quadricolor*. Cophylogenetic analysis using PACo and Random TaPas recovered highly significant congruent signal between host anemones and endosymbiotic Symbiodiniaceae. These patterns of association are indicative of high specificity and coevolutionary dynamics between symbiotic partners. Our results from cophylogenetic signal indicate that past evolutionary history has contributed to shaping the current interactions between giant sea anemones and Symbiodiniaceae strain. However, the observed pattern may emerge through multiple non-exclusive mechanisms [52]. This includes host-associated diversification of Symbiodiniaceae lineages, preferential host shifts among phylogenetically related hosts, ecological filtering driven by conserved host traits, or parallel responses of host and symbiont to shared environmental conditions. Distinguishing among these scenarios will require integrating cophylogenetic analyses with additional ecological, functional, evolutionary, and biogeographic data.

These patterns of Symbiodiniaceae host specificity are unsurprising as Japanese giant sea anemones vertically transmit algal symbionts. Giant sea anemone eggs contain bundles of Symbiodiniaceae from the parent anemones, or reproduction can occur asexually via longitudinal fission [53,54]. Vertical transmission, when the offspring inherits the same symbiont as the parent, is often associated with host specificity and co-evolution [24,55]. This results in high fidelity between symbiotic partners, with numerous benefits such as increased stability, heightened bleaching resilience, and specialized communication and metabolic processes [56,57]. The strong clustering of Symbiodiniaceae lineages according to anemones in this study is consistent with vertically transmitting symbioses and further supports the presence of host-specific symbiont strains. Such fidelity may provide the evolutionary context in which physiological integration between anemones and Symbiodiniaceae can arise and be maintained.

### Cryptic *Entacmaea* lineages harbor distinct *Cladocopium* species

Three-dimensional reconstructions of Symbiodiniaceae associated with lineages A and D of *Entacmaea quadricolor* reveal consistent differences in cell size and internal organization, particularly in the relative proportions of chloroplasts and lipid droplets. Although the limited number of reconstructed cells precludes robust statistical testing, the magnitude and consistency of these differences suggest distinct cellular strategies in carbon storage and photosynthetic investment, rather than environmental or developmental effects. Volumetric comparisons among Symbiodiniaceae remain relatively rare, although diagnostic characteristics such as chloroplast structure and volume have previously been used in some species descriptions [58,59]. Combined with the signatures of positive selection observed in several genes and the presence of amino acid substitutions within conserved functional domains, the morphological differences observed here suggest that divergence among Symbiodiniaceae associated with cryptic *Entacmaea* lineages is not limited to gene sequences but also extends to cellular organization and likely physiology and ecology.

Further identification of genes under positive selection in Symbiodiniaceae associated with *Entacmaea* lineages suggests different adaptive trajectories. Though the function of Symbiodiniaceae proteins remains poorly characterized [60], several of the identified genes are involved in functionally important processes within the symbiosome. Given the continual mutual exchange of carbon, nitrogen, and metabolites between host and symbiont, differences in functional categories of genes may reflect lineage-specific selective pressures acting on nutrient acquisition and cellular morphology. C_EquadD contains several positively selected genes involved in carbon flux, energy metabolism and metabolite transport. This observation is concordant with the ultrastructural data, showing an increased proportional volume of chloroplasts in C_EquadD than C_EquadA. Conversely, C_EquadA has more genes associated with amino acid and lipid metabolism, while containing a larger volume of lipid droplets. These differences may reflect levels of nitrogen made accessible to Symbiodiniaceae by their host anemone [6]. Nutrient availability may have consequential effects on Symbiodiniaceae morphology; for instance, environments low in nitrogen may cause a reduction in chloroplasts and an increase in lipid droplet volume, whereas abundant nitrogen access may have the opposite effect [61,62]. These interpretations remain necessarily speculative due to the limited knowledge surrounding Symbiodiniaceae gene function and morphology. The positively selected genes do not directly explain the observed ultrastructural differences, but the correspondence between functional categories under selection and lineage-specific cellular phenotypes suggest a coordinated physiological divergence. These parallel patterns suggest genomic and cellular differentiation between C_EquadA and C_EquadD.

These results become even more interesting when viewed in the broader context of the three-way symbiosis between anemonefish, sea anemones, and Symbiodiniaceae. *Entacmeaea* lineages A, B and C are preferentially associated with *Amphiprion clarkii*, whereas lineage D is with *A. frenatus*. Remarkably, this ecological structuring is mirrored by the algal symbionts, with each cryptic anemone lineage hosting a distinct lineage of *Cladocopium*. Thus, phylogenomic, ecological, and cellular datasets all converge toward the existence of distinct biological entities associated with these cryptic *Entacmaea* lineages. Interestingly, phylogenies indicate Symbiodiniaceae belonging to *E. quadricolor* lineage A as forming a distinct clade relative to the symbionts associated with other *Entacmaea* lineages. The presence of specific symbionts further supports the existence of cryptic *Entacmaea* lineages and suggests that Symbiodiniaceae composition is more closely associated with host identity than resident anemonefish species. With this, Okinawan *E. quadricolor* have two cooccurring symbiotic patterns, each centered around the host anemone (**Supp. Figure 3).** With their high specificity, we believe resident Symbiodiniaceae may serve as an additional line of evidence for determining anemone species’ identity. Altogether, our results suggest that diversification of this iconic symbiotic system involved not only anemonefish and their host anemones, but also their algal partners, revealing a previously hidden dimension of co-evolution.

Our preliminary genomic analyses suggest that the cryptic lineages of *Entacmaea* may themselves correspond to distinct species. We therefore have a powerful experimental system combining distinct lineages of anemone, anemonefish, and Symbiodiniaceae. Such a system offers a unique opportunity to dissect the mechanisms underlying partner specificity within this tripartite symbiosis. For example, it should allow testing whether Symbiodiniaceae contributes directly to the cues used by anemonefishes to recognize their host anemones, exploring how the metabolism of the three partners is interconnected, or identifying the molecular mechanisms responsible for the fidelity observed between specific fish, anemone, and algal lineages. More generally, this system may provide a particularly useful framework to understand how interactions among symbiotic partners contribute to their ecological specialization, co-evolution, and ultimately speciation.

## Materials and Methods

### Samples

Fifty-one giant sea anemone tentacles belonging to 6 species (*Heteractis aurora*, *H. magnifica Radianthus. crispa*, *Stichodactyla gigantea*, and *S. mertensii* and all four *E. quadricolor* cryptic lineages.) were collected across the Ryukyu Islands in Japan. *E. quadricolor* anemones were additionally collected from Ogasawara (Lineage B) and Shikine (Lineage C), near the northern limits of their distribution. These samples were previously used in Kashimoto et al. 2024 for investigation into giant sea anemone phylogeny, though this study filtered Symbiodiniaceae transcripts from host cnidarian for downstream analysis.

Additional sea anemones were collected from Okinawa and kept in aquaria at the OIST Marine Science Station for supplemental anemone tentacle collection for structural assessment.

Note that the taxonomy of clownfish-hosting sea anemones is currently undergoing substantial revision as molecular phylogenetic studies continue to reveal inconsistencies in traditional genus-level classifications. Although *H. magnifica* does not belong within *Heteractis* based on recent molecular phylogenies [12,13], we follow the current taxonomic consensus pending formal revision of the group.

### Sequencing

This study shares samples with Kashimoto et al. 2024 [21], further information on anemones, collection, and sequencing can be found there and Kashimoto et al. 2022 [13]. Briefly, giant sea anemone tentacles were collected across Japan. These samples were homogenized in 700 µL of Trizol (Thermo Fisher Scientific, USA), with RNA extracted according to manufacturer’s protocol. RNA quality was assessed via gel electrophoresis, and high quality samples underwent library construction through TruSeq Stranded mRNA kit (Illumina, USA). Samples were sequenced using NovaSeq6000 instrument, yielding an average of 65.3 million paired-end 150 base pair reads. **Supplementary Table 2** contains sample collection location information, SRR accession number, and BioSample number.

*De novo* transcriptomes containing both anemone and Symbiodiniaceae sequences were assembled using Trinity (ver. 2.12.0) [63]. Post assembly, transcripts were compared against custom blastn databases containing the transcriptomes of the seven Japanese sea anemones and the following Symbiodiniaceae species: *Symbiodinium microadriaticum* [64], *Breviolum minutum* [64], *Cladocopium goreaui* [65], and *Durusdinium trenchii* [66]. Only high-quality transcripts belonging to only host or symbiont were used. The Symbiodiniaceae transcripts constituted 25.4-46.5% of the complete assemblies, which correspond with previous studies on anemone tissue RNA sequencing [13,21]. After additional cleaning via cd-hit (ver. 2016-0304) [67] and ensuring complete removal of cnidarian transcripts, marker sequences were compared to ITS [38,39] and LSU [27,40] databases via blastn, with an e-value cutoff of 1e-20 (ver. 2.7.1+) [45]. Sequences were aligned via mafft (ver 7.505-1) [68] and visualized using via Geneious Prime (ver. 2025.1.3) (https://www.geneious.com/) for inspecting and cleaning sequences. Partial 5.8 and 28S sequences were trimmed from ITS2 sequences for full analysis of only the ITS2 marker [39]. Phylogenetic trees were generated through iqtree2 (ver 2.0.7+dfsg-1+b2) [69] and visualized with figtree (ver. 1.4.4-6) (accessible from: https://tree.bio.ed.ac.uk/software/figtree/).

### Phylotranscriptomics

Protein sequences were generated from transcriptomes using Transdecoder, using the setting – single-best-only (ver 5.7.1-2). Orthologous proteins between samples were clustered via OrthoFinder (ver. 2.0.4) [70] to determine the relationship between Symbiodiniaceae.

OrthoFinder was used to assign orthologous gene groups implementing Multiple Sequence Alignment (MSA). A treefile generated by the program creates a complete multi-locus tree, along with the file Orthogroups_for_concatenated alignment, containing the orthogroups used to create the tree. Single copy genes identified through OrthoFinder were concatenated using AMAS [71] [Borowiec 2016]. Tree constructions were performed using the Maximum Likelihood approach. Phylotranscriptomic analysis included datasets from *Symbiodinium microadriaticum* [64], *Breviolum minutum* [64], *Cladocopium goreaui* [65], and *Durusdinium trenchii* [66]. For easier visualization, trees were pruned to just contain the anemone-derived samples and the *C. goreaui* outgroup. The pipeline for phylotranscriptomics can be found here: https://github.com/ethanrickards/Symbiodiniaceae_phylotranscriptomics/tree/main

### Cophylogenetics

Two complementary methods were used to assess cophylogenetic signal between host (sea anemones) and symbiont (Symbiodiniaceae) evolutionary histories. Firstly, the global assessment of cophylogenetic signal was performed using the R-package paco [72] implementing the Procrustean Approach to Cophylogeny (PACo) [46]. Host and symbiont phylogenies were converted into cophenetic distance matrices and projected into Euclidean space using principal coordinates analysis. We applied a Cailliez correction and phylogenetic congruence was quantified through an asymmetric Procrustes superimposition, assuming that symbiont diversification is conditioned by host evolutionary history. Statistical significance of the Procrustes sum of squared residuals (SS) was assessed using 9,999 permutations of the association matrix under the r0 null model. The resulting SS statistic and associated permutation-based P-value were used to evaluate the strength of cophylogenetic signal. Secondly, we applied Random Tanglegram Partitions (Random TaPas) [47] using the R-package Rtapas [73] to identify the host–symbiont associations contributing most strongly to the observed cophylogenetic signal. Random TaPas repeatedly generates random partial tanglegrams of fixed size and evaluates their phylogenetic congruence using a global-fit statistic. We used PACo as the underlying congruence metric and generated 10,000 random partial tanglegrams, each containing 20% of the total number of observed associations. For each iteration, the Procrustes goodness-of-fit statistic was calculated, and the top 1% most congruent partial tanglegrams were retained. The contribution of individual associations to the overall cophylogenetic signal was quantified using weighted frequencies (wFq), which measure how often each association occurred among the most congruent subsets of interactions. Positive wFq values indicate associations contributing disproportionately to phylogenetic congruence, whereas negative values indicate comparatively weak contributions. The tangle_gram() function was used to provide a graphical summary of the Random TaPas analysis by mapping the weighted frequencies (wFq) of individual host–symbiont associations onto the tanglegram.

### Positively Selected Gene Comparison

Protein sequences of the Symbiodiniaceae belonging to *E. quadricolor* lineages were directly compared using Posigene [48]. Both C-EquadA and C-EquadD were selected as anchor species to detect positive selection in the specified lineage against multiple species. The Symbiodiniaceae belonging to the opposing Symbiodiniaceae lineage, *Cladocopium goreaui*, and *Symbiodinium microadriaticum* were all used as non-high-scoring BLAST references. After candidate genes were selected, the protein functionality was predicted using interproscan (ver. 5.76-107.0) [74] and blastp [45] against reference datasets of nr, *Arabidopsis sp., Homo sapiens,* and Cnidaria [49]. Functionally important protein domains were identified via SMART Web portal [75].Each gene was then aligned using mafft [68] and amino acid sequence visually compared via Geneious Prime (ver. 2025.1.3). To calculate percent identity, positively selected genes were compared using BLASTn against databases containing transcriptomes of *E. quadricolor* D and A, along with *C. goreaui* and *D. trenchii*. The highest quality hit with complete coverage of each gene per sample was isolated and used to calculate percent identity.

### 3D imaging

3-4 tentacles were collected from *Entacmaea quadricolor* kept in aquaria for structural analysis. Sea anemone tentacles tips were collected with scissors and immediately fixed in 2.5% glutaraldehyde (Sigma-Aldrich, St. Louis, MO, USA) prepared in 0.1 M phosphate buffer (pH 7.4) for 2 hr at room temperature. Following primary fixation, samples were rinsed three times in 0.1 M phosphate buffer for 10 minutes each and post-fixed in a solution containing 1.5% potassium ferrocyanide (FeCN) and 2% osmium tetroxide (OsO_4_) in 100 mM phosphate-buffered saline (PBS) at pH 7.0.

Samples were rinsed five times in distilled water for 3 minutes and incubated in 1% thiocarbohydrazide for 20 minutes at room temperature. Afterwards, samples were rinsed five times in distilled water for three minutes and incubated in 2% osmium tetroxide for 30 minutes at room temperature. Excess staining solution was removed through five additional rinses in distilled water, then incubated overnight in 1% uranyl acetate at 4℃.

Samples were washed five times in distilled water prior to dehydration. Dehydration was carried out through a graded ethanol series, with five minutes at each of the following steps: 20%, 50%, 80%, 90%, and 100% ethanol. Samples were then incubated in a 1:1 mixture of acetone and ethanol for 30 minutes, followed by 100% acetone for an additional 30 minutes.

Resin infiltration conducted by incubating the samples in a 1:1 mixture of acetone and epoxy resin for 2 minutes. Afterwards, samples were transferred to 100% resin and incubated overnight at room temperature. Resin was replaced the next day with fresh 100% resin and samples were incubated for an additional 1 hour. Samples were transferred into embedding molds and polymerized at 60℃ for 72 hours.

### Serial block-face scanning microscopy (SBF-SEM)

For three-dimensional ultrastructural analysis, resin blocks were mounted onto aluminum pins using conductive silver epoxy (Electron Microscopy Sciences, #12642-14, Hatafield, PA 19440, USA) and trimmed to expose the cross-section of the tentacle. Serial block-face imaging was performed using a Thermo Fisher Scientific TeneoVS Electron Microscope (ThermoFisher Scientific, Waltham, Massachusetts, USA) equipped with an integrated ultramicrotome. The integrated ultramicrotome sequentially removed thin layers of the resin block while images were acquired from freshly exposed block surface. Imaging was conducted via backscattered electron detection at an accelerating voltage of 2 kV and beam current of 1.5 nA. Pixel size in the x-y plane was 6.132 nm, and thickness 0.06 µm, resulting voxel size was 6.132 x 6.132 x 60 nm.

Image stacks obtained from SBF-SEM were processed using Amira3D Software (version 2025.1). Image stacks were stitched using Bigstitcher (https://github.com/ECBSU/imaging-scripts/blob/main/volume_em/Bigstitcher_SBF_Teneo.ijm). Slices were aligned, contrast normalized and noise filtered were performed using Microscopy Image Browser (Belevich et al. 2016). Image slices were cropped to only contain individual cells. Manual segmentation was performed on chloroplast, mitochondria, nucleus, accumulation body, lipid droplet, starch body, and pyrenoid. Other non-labeled structures of the cell fell under “unlabeled cell structure”. Segmentation of criteria for each structure was defined based on morphological features. Volume was calculated directly from voxel-based reconstructions. Reconstruction volumes were visualized and rendered using Amira ver. 2025.1 and Dragonfly ver. 2025.1.

## Acknowledgments

We would like to thank Filip Husnik, Javier Tejeda Mora, and Yong Heng Phua for their guidance on three-dimensional Symbiodiniaceae reconstruction. We thank OIST Imaging Section for anemone tentacle section microscopy. We are grateful for the help and support provided by OIST SQC for sequencing services, along with the husbandry assistance performed by OIST Marine Animal Research Support.

## Author Contributions

E.R.: Phylogenetic analysis, 3D image construction and analysis, positive selection analysis, and manuscript preparation. S.T.-P. and M.H.: SBF-SEM imaging and manuscript preparation. S.M.: Molecular laboratory work. M.W.: technical advice and design. B.F.: Cophylogenetic analysis and manuscript preparation. K.K.: Transcriptome assembly and filtering. V.L.: Manuscript preparation.

## Supplementary Figures

**Supp. Figure 1.**
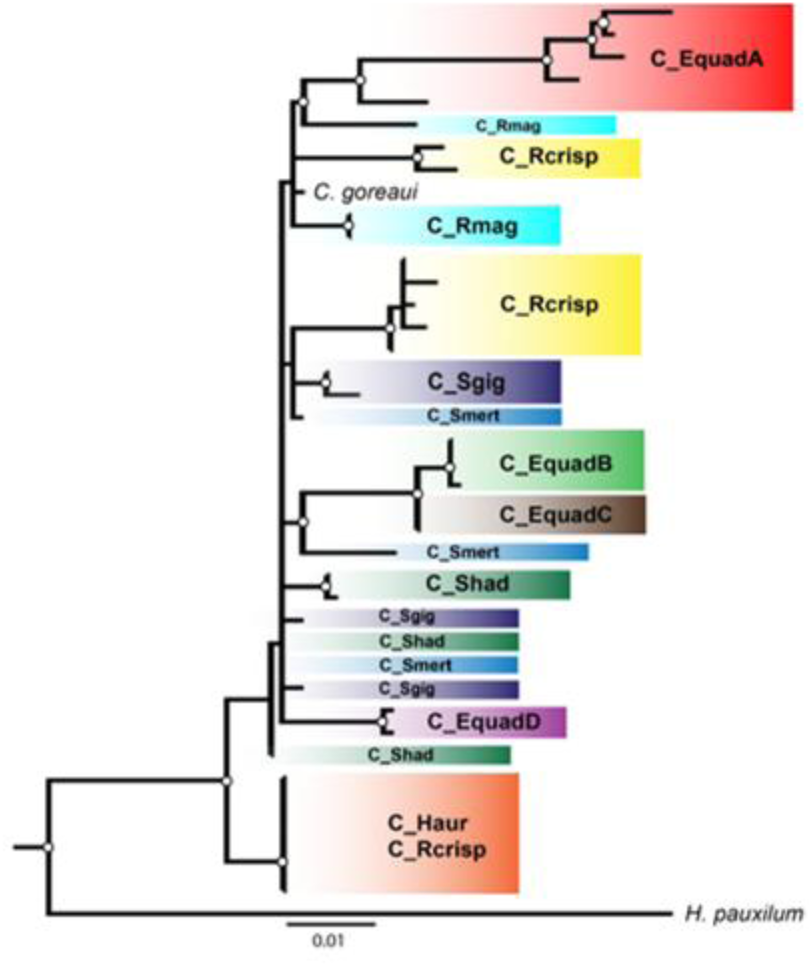
Phylogeny of Symbiodiniaceae through concatenation of ITS2 and LSU marker sequences. Marker sequences were determined through blastn comparison against ITS2 (Hume et al. 2019, Shi et al. 2021) and LSU (LaJeunesse et al. 2018, Butler et al. 2023) databases. Sequences were appended together to form an approximately 800-bp long sequence, aligned, and constructed via iqtree2. Samples missing either an ITS2 or LSU sequence were excluded from the tree.

**Supp. Figure 2.**
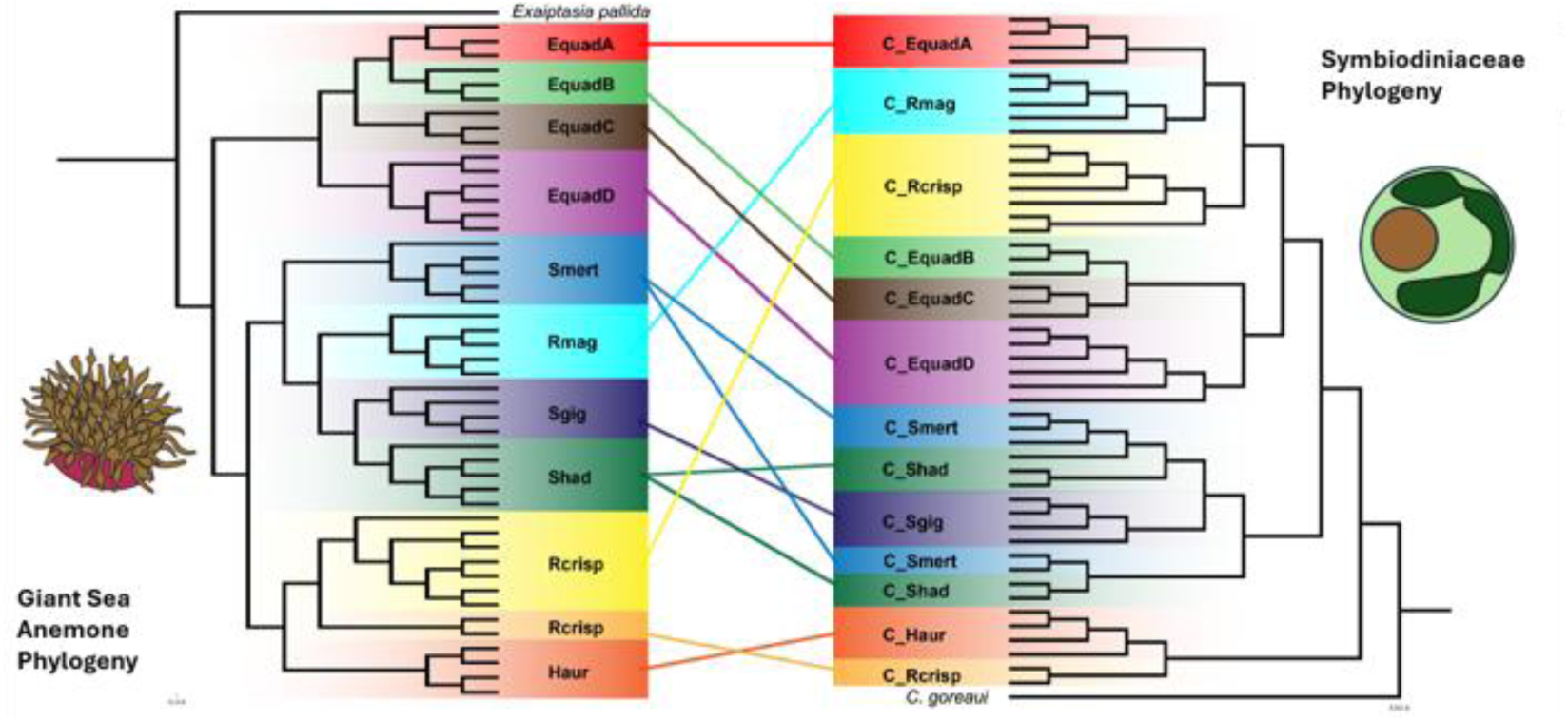
Comparison between Symbiodiniaceae and host giant sea anemone phylogenies. Maximum-likelihood phylogenetic trees were constructed using iqtree2. Each tip is color labeled according to host species, with color-coded lines between Symbiodiniaceae and their host anemone for easy comparison.

**Supp. Figure 3.**
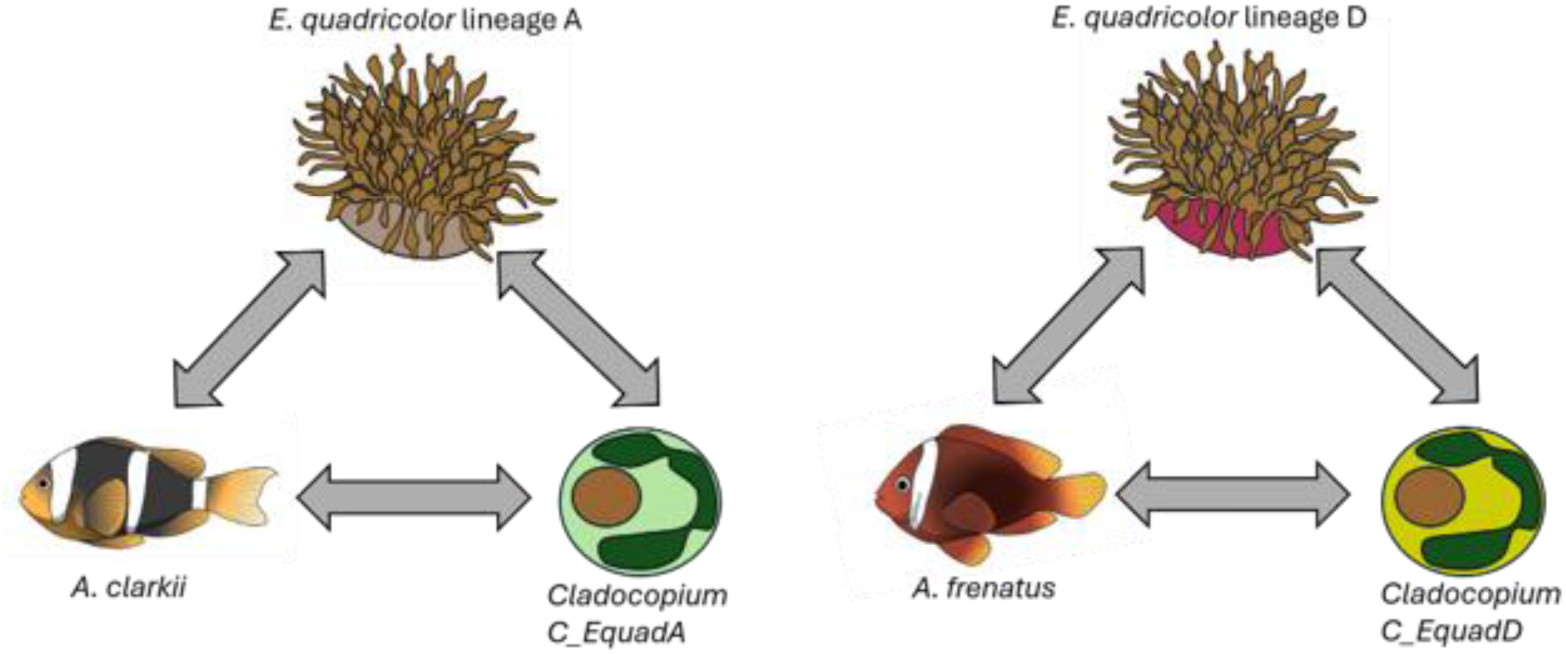
Two distinct symbiotic systems co-occurring in Okinawa. Each anemone–fish– Symbiodiniaceae system comprises a distinct combination of symbiotic partners.

